# Associations of maternal prenatal emotional complaints and cortisol with neonatal meconium microbiota: A cross-sectional study

**DOI:** 10.1101/2022.09.25.509368

**Authors:** Nadia Deflorin, Ulrike Ehlert, Rita T. Amiel Castro

## Abstract

The gut microbiome is a complex microbial ecosystem considered as a key modulator of human health and disease. Alterations in the diversity and relative abundances of the gut microbiome have been associated with a broad spectrum of medical conditions. Maternal emotional complaints during pregnancy can impact on offspring development by altering the maternal and the foetal gut microbiome. We aimed to investigate whether self-reported maternal anxiety, depressive symptoms, and distress as well as biological stress in late pregnancy alter the bacterial composition of the infant’s meconium.

**Methods:** A total of *N*=100 mother-infant pairs were included. Maternal emotional complaints were measured using standardised questionnaires (EPDS, PSS-10, STAI) at 34-36 weeks gestation and salivary cortisol was measured at 34-36 and 38 weeks gestation. Infant meconium samples were collected in the first five days postpartum and analysed using 16S rRNA amplicon sequencing.

**Results:** Correlations showed that lower alpha diversity of the meconium microbiome was significantly associated with increased maternal prenatal depressive symptoms in late gestation (τ = -0.149, *p* = 0.041). Increased cortisol AUCg at T2 was significantly related to higher beta diversity of the meconium samples (Pr(>F) = 0.003*). *Proteobacteria* was the most abundant phylum and was associated with maternal cortisol total decline. No other associations were found.

**Conclusions:** Maternal prenatal depressive symptoms are associated with infant faecal microbiota alpha diversity, whereas maternal cortisol AUCg is linked to increased beta diversity and total decline related to increased *Proteobacteria*. Future studies are warranted to understand how these microbiota community alterations are linked to child health outcomes.

**Highlights:**

- Maternal prenatal depressive symptoms alter the meconium microbiome.
- Increased biological stress during pregnancy alters the beta diversity of the meconium microbiome.
- *Proteobacteria* is the most abundant phylum in the meconium samples.

## 1. Introduction

Pregnancy is a significant life event, as the transition to motherhood is associated with major challenges in the psychological, biological and social domains [1, 2]. Indeed, many women report emotional complaints (e.g. stress, anxiety, depression) during pregnancy, with the prevalence of prenatal depression ranging from 5-25% [3–5] and the prevalence of prenatal anxiety estimated to be up to 21% [6]. Experiencing depression, anxiety, or stress during pregnancy can result in a range of altered neurodevelopmental outcomes in the offspring, including cognitive, developmental, and mental health difficulties from childhood into adulthood [7]. Similarly, prenatal depressive and anxiety symptoms have been linked to intrauterine growth restriction, prematurity, and low birth weight [8]. While some of the biological mechanisms underlying these associations are beginning to be understood, others remain unexplored [9].

One possible mechanism that has recently gained more attention is the gut microbiome [10]. The gut microbiome is a complex microbial ecosystem that is found on the intestinal mucosal surfaces and is considered as a key modulator of human health and disease. The microbiome-gut-brain axis refers to the bidirectional interaction between the gastrointestinal tract (GI), the intestinal microbiome, and the central nervous system [11]. Findings from animal models suggest a link between the intestinal microbial community and brain development [12, 13], while in humans, gut microbiome alterations have additionally been linked to the host’s health [14]. In particular, changes in the diversity and relative abundances in the gut microbiome have been associated with a broad spectrum of medical conditions including neurobehavioural and affective disorders [14–17]. Depression and anxiety can alter gut motility, which in turn may change the composition and stability of the gut microbiome, colon physiology and morphology [18–21]. In response to stress, the hypothalamic-pituitary-adrenal axis is stimulated, producing, inter alia, the stress hormone cortisol [9, 22]. Studies with pregnant rats revealed that higher maternal cortisol levels were associated with lower bacterial population in the offspring [23]. In humans, Zijlmans and colleagues [9] examined faecal samples from infants and reported that elevated prenatal stress experience and salivary cortisol concentrations were significantly related to higher relative abundances of pathogenic Proteobacterial groups and lower relative abundances of lactic acid bacteria in the infant’s gut microbiome. The diversity and composition of the gut microbiome modulates the activity of GI-specialised enteroendocrine cells, which release signalling molecules that have important endocrine and metabolic functions [24–27]. Therefore, in view of the extensive microbiome-host interactions, early gut bacterial colonisation is critical for the modulation of health and disease [28, 29].

Although the exact timing of the establishment of the infant gut microbiome remains unknown, findings from different research groups [9, 30, 31] support the notion of an intrauterine colonisation of the foetal intestinal gut. A number of studies have identified the presence of commensal bacteria in the placenta, in amniotic fluid, as well as in the uterus [32– 37], with potential bacterial translocation from the maternal intestine to the bloodstream [38– 40]. Studies in mice [41] and monkeys [42] have shown that maternal prenatal stress can affect the maternal intestinal microbiome and can subsequently expose the foetus to altered populations of microbes [41]. These alterations occur through different microbiome-gut-brain axis communication pathways, including the immune system, neurotransmitters, metabolites, as well as vagal nerve activation [15]. However, only a small number of human studies to date have explored this topic during pregnancy. A recent study including 410 mother-infant pairs examined the influence of prenatal emotional complaints on neurodevelopmental problems in the offspring [43]. The findings revealed that prenatal emotional complaints were associated with alterations in both the meconium (first stool discharge of newborns) microbiome and neurodevelopment at 24 months of age. Hu and colleagues (2019) [30] investigated whether maternal mental health was associated with the meconium microbiome of 75 newborns, and found an association between higher levels of pregnancy-specific anxiety and changes in bacterial composition. Finally, in a study with mother and infants (84 mothers and 101 newborns), Naudé and colleagues (2020) [31] reported that maternal lifetime intimate partner violence and prenatal psychological distress were associated with altered bacterial profiles in both infant and mother faeces.

Despite some prior research in this area, evidence is still lacking regarding the influence of maternal prenatal emotional complaints on the composition of the neonate’s intestinal microbiome measured in meconium [29]. Considering that self-reported distress, depressive symptoms, and anxiety are common in the perinatal period [5, 6, 44] and may influence early foetal gut colonisation [15, 41, 42, 45], further research is warranted. Therefore, we aimed to investigate whether maternal prenatal anxiety, depressive symptoms, and self-reported as well as biological distress alter the bacterial composition of the infant’s meconium. To examine this complex interplay, the present study incorporated a wide variety of recommendations in order to achieve the highest data quality. Multiple confounding factors were investigated, storage recommendations were carefully followed, both psychological and biological indices of stress were employed, and information relevant to the sample collection was obtained [46–48]. In line with the study’s aim, we hypothesised that maternal prenatal biological stress, self-reported distress, depressive symptoms and anxiety would be associated with the diversity and composition of the neonatal meconium microbiome.

## 2. Materials and Methods

This is a prospective cross-sectional study assessing perinatal women and their children (*N* = 100 mother-infant pairs). The study was approved by the Cantonal Ethics Committee (KEK-ZH-No. 2020-01928) and was conducted in accordance with the principles of the Declaration of Helsinki.

Participants were recruited in Switzerland from January 2021 to January 2022 through the study’s website and information leaflets that were handed to patients receiving prenatal medical care at maternity hospitals and gynaecological practices in the Swiss cantons of Zurich and Aargau. Pregnant women were also recruited through various mailing lists, online platforms, and social media (e.g. Facebook). Inclusion criteria for the present study were a) self-reported physically healthy women, b) currently being in the 3^rd^ gestational trimester, b) age between 18 and 45 years, c) fluency in German, and d) being of European origin. Exclusion criteria were: a multifoetal gestation, pregnancy due to assisted reproductive technology, medical complications and/or surgical interventions that might have affected ovarian functions prior to pregnancy (e.g. polycystic ovary syndrome), and a planned Caesarean section. Furthermore, women currently taking hormones, psychotropic substances, drugs (e.g. cocaine, marijuana), or antibiotics were excluded.

Interested pregnant women were screened for eligibility using an online questionnaire and a face-to-face interview. In the interview, the study procedure was explained, the inclusion and exclusion criteria were verified, and the Structured Clinical Interview for DSM-5 Clinician Version (SCID-5-CV) was administered by the first author. Eligible women who wished to participate and met the inclusion criteria (*N* = 119) received study information and consent forms via postal mail and were invited to an appointment between the 34^th^ and 36^th^ week of gestation. Prior to the appointment (T1), participants were asked to read the study information thoroughly and sign the consent form. This appointment took place either as a home visit or online via videotelephony, depending on the COVID-19 pandemic-related restrictions in place at the time of participation. At T1, the study procedure was explained and anthropometric measurements and blood pressure were recorded. Participants were then instructed on how to take the saliva samples and how to collect the neonate’s meconium. These instructions were provided both verbally and in writing, as the participants were required to carry out the saliva sampling and meconium collection independently during the course of the study. To facilitate meconium collection, participants received a faecal and saliva collection kits. Finally, participants were asked to complete online questionnaires in order to provide sociodemographic information and psychological and behavioural data. When the appointment was held online, the same procedure was applied and the anthropometric measures and blood pressure were taken from participants’ maternity records.

For the two consecutive days after T1, participants were asked to take saliva samples immediately after awakening, 30 min and 45 min after awakening, and before going to bed, and also to complete a brief questionnaire on perceived distress and sleep quality. A second saliva sampling, following the same protocol, was conducted on the first two days of the 38^th^ week of pregnancy (T2). In the postpartum phase, once the newborn had excreted the meconium, mothers were asked to collect it (e.g. from a diaper or birthing room) and to store it at room temperature in an appropriate vial provided with the faecal collection kit. The final study appointment (T3) was conducted between 8:00 and 8:30 in the morning as a home or clinic visit within the first 120 hours after birth, and preferably on the fourth day postpartum. At T3, we clarified any questions mothers might have regarding the study and took the meconium and saliva samples away for analysis. Additionally, we asked the women about their baby and birth characteristics.

### 2.1. Instruments

#### 2.1.1. Maternal emotional complaints

At T1, psychometrically validated questionnaires were used to assess prenatal emotional complaints, including depressive symptoms, anxiety symptoms, and perceived stress.

Maternal depressive symptoms were assessed using the German version of the *Edinburgh Postnatal Depression Scale* (EPDS) [49], a widely used questionnaire designed to assess depressive symptoms in pregnant and postpartum women. The scale consists of 10 items rated on a four-point Likert scale (0 = “*yes, most of the time*” to 3 = “*no, never*”) referring to the last seven days, with the total score ranging from 0 to 30. In the present sample, Cronbach’s alpha lay at 0.798. The total EPDS score was used as a continuous variable in the analyses.

The German version of the *State-Trait Anxiety Inventory* was administered to measure trait anxiety (STAI-T) and state anxiety (STAI-S) [50, 51]. Trait anxiety refers to the general tendency to experience worries, fears, and anxieties across various situations [52]. The STAI-T scale contains 20 items rated on a four-point Likert scale (e.g., from 1 = “*almost never*” to 4 = “*almost always*”), with higher scores indicating higher trait anxiety. Cronbach’s alpha for this scale in the present sample lay at 0.891. State anxiety, by contrast, refers to the degree of anxiety experienced in a current situation [50]. The STAI-S scale likewise consists of 20 items rated on a four-point Likert scale (1 = “*not at all*” to 4 = “*very much*”), with higher values indicating higher state anxiety [53]. Cronbach’s alpha in the present sample lay at 0.897. The total score was used as a continuous variable in the analyses.

The *Perceived Stress Scale (PSS-10)* is a well-established 10-item self-report scale measuring the extent to which the respondent considers events during the past four weeks to be unpredictable, uncontrollable, and stressful, with items rated on a five-point Likert scale (e.g., 0 = “*never*” to 4 = “*very often*”) [54]. Higher scores indicate a higher level of perceived stress. The German version of the scale was used in the present study [55] and Cronbach’s alpha in our sample lay at 0.828. The total PSS-10 score was used as a continuous variable in the analyses.

#### 2.1.2. Maternal prenatal salivary cortisol

In total, each participant collected 16 saliva samples under standardised conditions for the assessment of salivary cortisol concentrations. Participants were asked to avoid exercise, alcohol, coffee, black tea, and chewing gum on the day before saliva sampling. Moreover, they were requested not to smoke, eat, or brush their teeth during the morning of saliva collection. The samples were collected using the passive drool method in SaliCap sampling tubes (2 mL) (SaliCaps, IBL international GmBH, Hamburg, Germany). The women were instructed to collect saliva immediately after awakening, 30 min and 45 min after awakening, and before going to bed, and to store the samples in their home freezers until T3. At T3, samples were transported in a cooling box and stored at -20 degrees in the biochemical laboratory at the Department of Clinical Psychology and Psychotherapy, University of Zurich. For the subsequent analyses, samples were thawed, centrifuged, and biochemically analysed. Analysis was performed using enzyme-linked immunosorbent assay kits (ELISA; IBL international GmBH, Hamburg, Germany). Individual values outside the range of the kits were excluded from the analyses (*N* = 2).

### 2.2. Neonatal meconium microbiome sample processing

Mothers were asked to collect the newborn’s meconium immediately after its excretion using a sterile faecal collection kit (OMNIgene GUT sample collection kit, OM-200) including a sterile spoon. The average time between meconium excretion and collection was 85.03 minutes (*SD* = 248.619). After collection, participants were required to close the vial tightly and shake it until the biomass was mixed with a liquid solution. The majority of meconium samples were collected from the diaper (66%), with some being directly collected from the neonate’s anus (11%) and some from both diaper and anus (6%). The remaining samples (17%) were collected from a towel (6%), the mother’s abdomen (3%), the floor (1%), a birthing chair (1%), or other surfaces (6%). After collection, participants were asked to store the samples at room temperature protected from sunlight. Members of the research team then collected the samples at T3 and transported them in a cooling box to the biochemical laboratory of the University of Zurich, where samples were stored at - 80 degrees until shipment. Samples were shipped to the Clinical Microbiomics laboratory (Copenhagen, Denmark) for DNA sequencing, using dry ice at -80 degrees.

### 2.3. Meconium 16S ribosomal RNA (rRNA) sequencing and quality control

At the Clinical Microbiomics laboratory, a NucleoSpin 96 Stool (Macherey-Nagel) kit was used to extract bacterial DNA from ∼200μL aliquots of the meconium samples. Aliquots were bead-beaten horizontally on a Vortex-Genie 2 at 2700 rpm for 5 minutes. Negative and positive (mock) controls were included in the analysis. Products from nested PCR were pooled based on band intensity, and the resulting library was purified with magnetic beads. A fluorometric measurement of the DNA concentration of the pooled libraries was performed. Sequencing was conducted on an Illumina MiSeq desktop sequencer using the MiSeq Reagent Kit V3 (Illumina) for 2x 300 bp paired-end sequencing. PCR was performed with the forward primer S-D-Bact-0341-b-S-17 and reverse primer S-D-Bact-0785-a-A-21 [56] with Illumina adapters attached. These are universal bacterial 16S rDNA primers which target the V3-V4 region. The following PCR program was used: 98 °C for 30 sec, 29x (98° C for 10 s, 55 °C for 20 s, 72 °C for 20 s), 72°C for 5 min. Amplification was verified by running the products on an agarose gel. Indices were added in a subsequent PCR using an Illumina Nextera kit with the following PCR program: 98 °C for 30 sec, 8x (98° C for 10 s, 55 °C for 20 s, 72 °C for 20 s), 72 °C for 5 min. Attachment of indices was verified by running the products on an agarose gel. All samples were rarefied to a fixed number of reads in order to account for variation in detection ability due to differences in sequencing depth. Abundance matrices and rarefied count were obtained by random sampling of reads without replacement, with a target number of 4063 read counts. Samples with lower read counts were excluded (*N* = 5). After the first sequencing run, 15 meconium samples had insufficient reads, of which 10 were successfully reextracted and resequenced. For the remaining 5 meconium samples, there was no remaining sample material to obtain good sequencing data.

### 2.4. Statistical analysis

Statistical analyses were performed using the IBM Statistical Package for the Social Sciences (SPSS Version 28 for Windows) and the software program R, using the adonis2 function from the vegan R package. An a priori power analysis was conducted using G*Power version 3.1.9.7 [57] to determine the minimum sample size required to test the study hypothesis. The Shapiro-Wilk test was applied to test for normal distribution. Non-parametric tests were used for non-normally distributed data.

Using a customised pipeline based on dada2, the meconium microbiome composition was summarised into an ASV abundance table [58]. Taxonomic classification of the discovered ASVs was performed in two steps. Initially, ASV sequences were compared with full-length 16S sequences in an internal reference database (CM_16S_27Fto1492R_v1.0.0) using a naive Bayesian classifier. In the second annotation step, taxonomic assignments were optimised using the exact sequence identity percentages between the detected ASVs and the reference amplicons in an internal V3V4 amplicon database (CM_16S_341Fto785R_v1.0.0.rds). Second, we examined the diversity of the meconium microbiome communities within and between samples. Therefore, bacterial alpha diversity was determined by the number of detected entities (richness) and by Shannon index, calculated from rarefied ASV abundances. Bacterial beta diversity was computed as Bray-Curtis dissimilarity as well as weighted UniFrac distances using rarefied data. Associative analyses with alpha diversity and taxon abundance as dependent variables and psychological variables as independent variables were conducted using Kendall rank correlations. The associative analyses computed for beta diversity - with the masses of Bray Curtis dissimilarity as well as weighted UniFrac distances as dependent variables and psychological variables as independent variables - were performed using a distance-based redundancy analysis (dbRDA) test conducted with 1000 permutations. For associations with microbial taxon abundances, we used unrefined data, for which *N* = 100 meconium samples were considered. Conversely, for the associations with alpha and beta diversity parameters, which were calculated based on rarefied data, we considered *N* = 95 meconium samples. When performing the statistical analyses, the Benjamini–Hochberg (BH) method was used to control for the false discovery rate (FDR) at a level of 10 % [59].

For the analysis of maternal salivary cortisol concentrations, both the mean value of the area under the curve with respect to the ground (AUCg) and the mean value of the total decline were calculated by taking the average of the cortisol concentrations on the two consecutive sampling days for these two specific parameters. This procedure was applied separately at T1 and T2. The total decline was calculated as follows: Δ in the morning - Δ during the day, with Δ in the morning corresponding to the highest morning value - baseline morning value and Δ during the day corresponding to the highest morning value - the evening value. To calculate AUCg, we used the trapezoid formula, which is routinely used to incorporate multiple time points [60]. All cortisol results were reported in nmol/l. Associative analyses between psychological markers of T1 (EPDS, PSS-10, STAI-S) and both the AUCg and the total decline were conducted using Spearman’s rank correlation. The same test was performed to calculate the correlations of the cortisol concentrations (AUCg and total decline) between the two time points.

#### 2.4.1. Covariates

The following covariates were identified from the literature [30, 31, 61–64]: age, parity, maternal BMI, newborn length and weight (treated as continuous variables), and education, mode of delivery and meconium collection site (coded as binary variables). To explore the possible influence of these covariates on the microbiome, permutational multivariate analysis of variance (PERMANOVA) and distance-based redundancy analysis (dbRDA) (similarly to the approach used by Falony et al., 2016 [65]) were performed. Only covariates that showed a significant association with the meconium microbiome were included in the analysis.

## 3. Results

### 3.1. Sample characteristics

Results of the a priori power analysis [57] indicated a required sample size of *N* = 84 to achieve 80% power to detect a medium effect (f = 0.15) at a significance level of α = .05. Accordingly, the obtained sample size of *N* = 100 is adequate to test the study hypothesis. Of the *N* = 119 participants who initially consented to participate, *n* = 12 were excluded due to medical complications and *n* = 1 due to a planned Caesarean section. A further *n =* 6 women dropped out of the study. The final sample thus comprised *N* = 100 pregnant women aged between 22 and 40 years, which is comparable to or larger than the samples in other studies in the field (e.g. Naudé et al. (2020) [31]).

Table 1 shows the descriptive characteristics of the sample. The participants’ median age was 32.50 years (IQR = 4.00) and the majority were from Switzerland (77%) or Germany (10%). Most participants had a high level of education, with 67% reporting a university degree or higher. For 54% of the study participants, this was their first pregnancy.

**Table 1.**
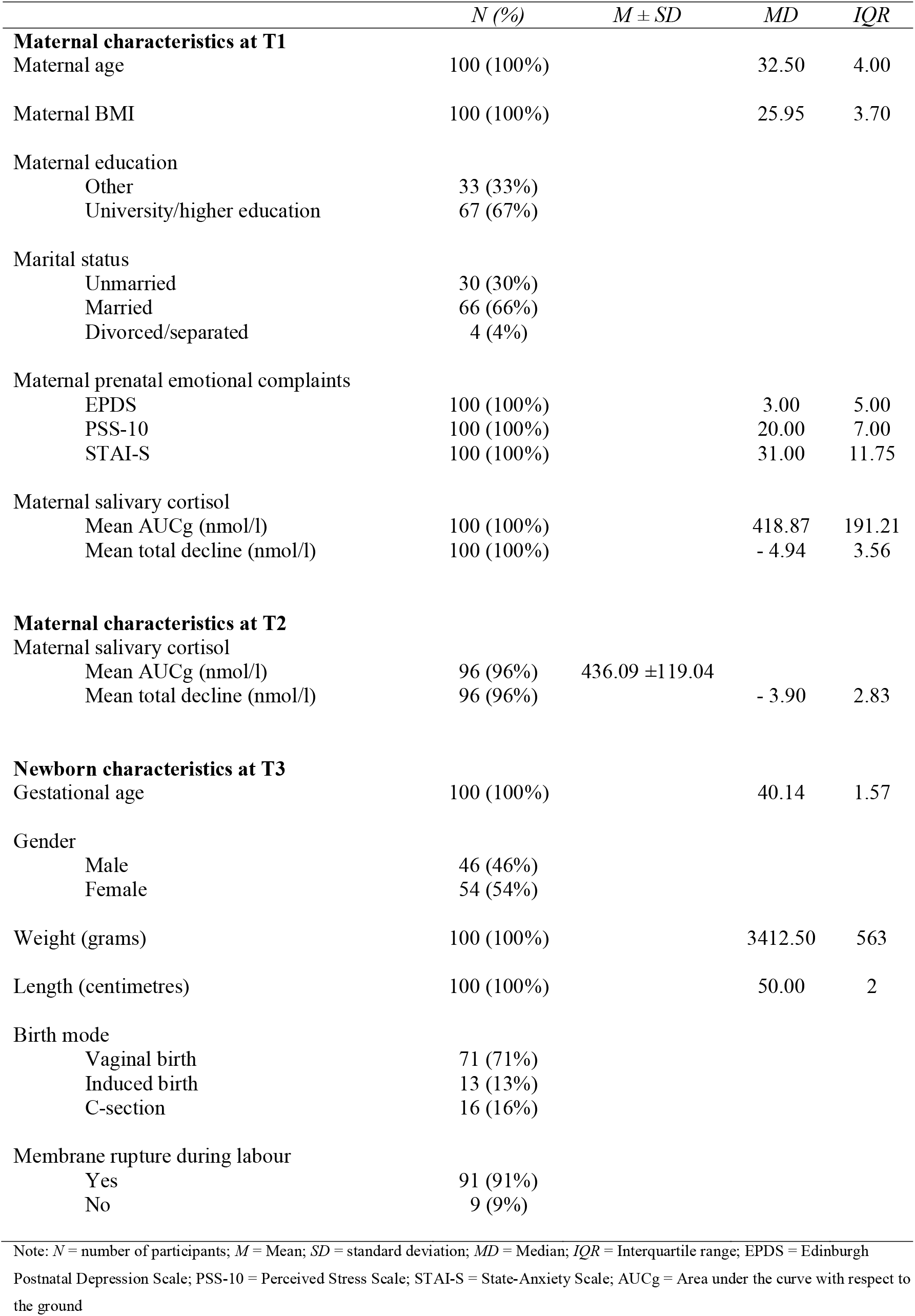
Demographic and obstetric outcomes of the study sample (*N* = 100)

| | $N$ (%) | $M \pm SD$ | $MD$ | $IQR$ |
| --- | --- | --- | --- | --- |
| **Maternal characteristics at T1** |  |  |  |  |
| Maternal age | 100 (100%) |  | 32.50 | 4.00 |
| Maternal BMI | 100 (100%) |  | 25.95 | 3.70 |
| Maternal education |  |  |  |  |
| Other | 33 (33%) |  |  |  |
| University/higher education | 67 (67%) |  |  |  |
| Marital status |  |  |  |  |
| Unmarried | 30 (30%) |  |  |  |
| Married | 66 (66%) |  |  |  |
| Divorced/separated | 4 (4%) |  |  |  |
| Maternal prenatal emotional complaints |  |  |  |  |
| EPDS | 100 (100%) |  | 3.00 | 5.00 |
| PSS-10 | 100 (100%) |  | 20.00 | 7.00 |
| STAI-S | 100 (100%) |  | 31.00 | 11.75 |
| Maternal salivary cortisol |  |  |  |  |
| Mean AUCg (nmol/l) | 100 (100%) |  | 418.87 | 191.21 |
| Mean total decline (nmol/l) | 100 (100%) |  | - 4.94 | 3.56 |
| **Maternal characteristics at T2** |  |  |  |  |
| Maternal salivary cortisol |  |  |  |  |
| Mean AUCg (nmol/l) | 96 (96%) | 436.09 ± 119.04 | | |
| Mean total decline (nmol/l) | 96 (96%) |  | - 3.90 | 2.83 |
| **Newborn characteristics at T3** |  |  |  |  |
| Gestational age | 100 (100%) |  | 40.14 | 1.57 |
| Gender |  |  |  |  |
| Male | 46 (46%) |  |  |  |
| Female | 54 (54%) |  |  |  |
| Weight (grams) | 100 (100%) |  | 3412.50 | 563 |
| Length (centimetres) | 100 (100%) |  | 50.00 | 2 |
| Birth mode |  |  |  |  |
| Vaginal birth | 71 (71%) |  |  |  |
| Induced birth | 13 (13%) |  |  |  |
| C-section | 16 (16%) |  |  |  |
| Membrane rupture during labour |  |  |  |  |
| Yes | 91 (91%) |  |  |  |
| No | 9 (9%) |  |  |  |

Note: $N$ = number of participants; $M$ = Mean; $SD$ = standard deviation; $MD$ = Median; $IQR$ = Interquartile range; EPDS = Edinburgh Postnatal Depression Scale; PSS-10 = Perceived Stress Scale; STAI-S = State-Anxiety Scale; AUCg = Area under the curve with respect to the ground

### 3.2. Analysis of meconium microbiome

Following data pooling of the two sequencing runs, the 100 meconium samples had an average of 75.2 thousand (K) read pairs per sample and a minimum of 3.6 K read pairs. A mean of 32.9 K read pairs per sample could be mapped to the ASV catalogue, representing an average of 44.7 % of the high-quality reads. For a taxonomic overview, Figure 1 and Figure 2 show the relative abundance of bacterial taxa aggregated at the genus and family levels in the meconium. We found that the meconium microbiome was dominated by the genera Acinetobacter (39.11%) and Pseudomonas (21.36%) (Figure 1). At the family level (Figure 2), we found that most of the assigned meconium taxa were from Moraxellaceae (39.90%) and Enterobacteriaceae (10.14%).

**Figure 1.**
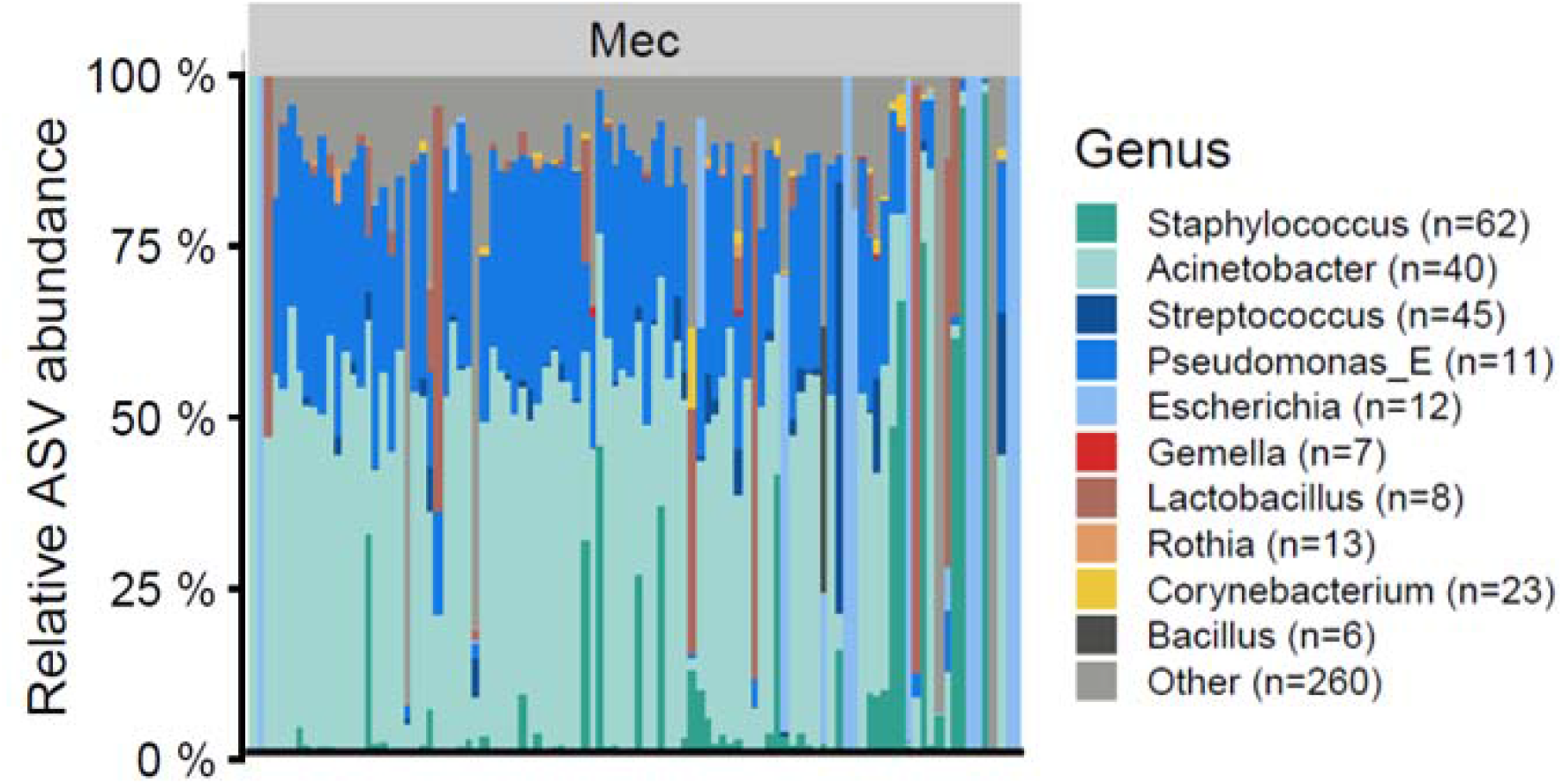
Taxonomic overview of the meconium microbiome at the genus level per sample for unrarefied data. The numbers in parentheses refer to the number of ASVs aggregated for a given taxon. Bar plots display the relative abundance of the top 10 taxa with the highest average abundance across all samples. Light grey (Other) indicates the total relative abundance of ASVs that are not in the top 10 most abundant taxa; Mec = Meconium.

**Figure 2.**
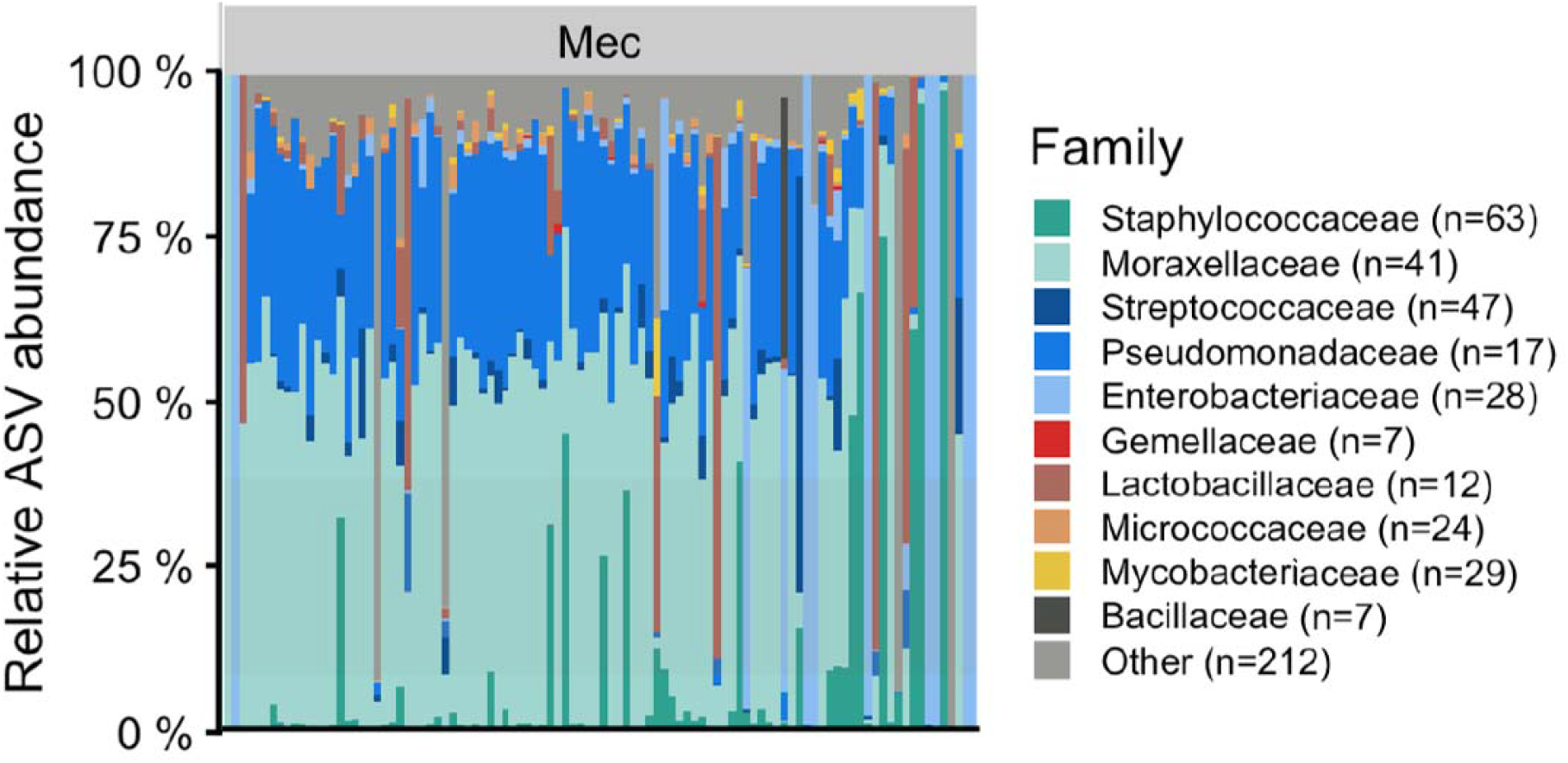
Taxonomic overview of the meconium microbiome at the family level per sample for unrarefied data. The numbers in parentheses refer to the number of ASVs aggregated for a given taxon. Bar plots display the relative abundance of the top 10 taxa with the highest average abundance across all samples. Light grey (Other) indicates the total relative abundance of ASVs that are not in the top 10 most abundant taxa; Mec = Meconium.

**Figure 3.**
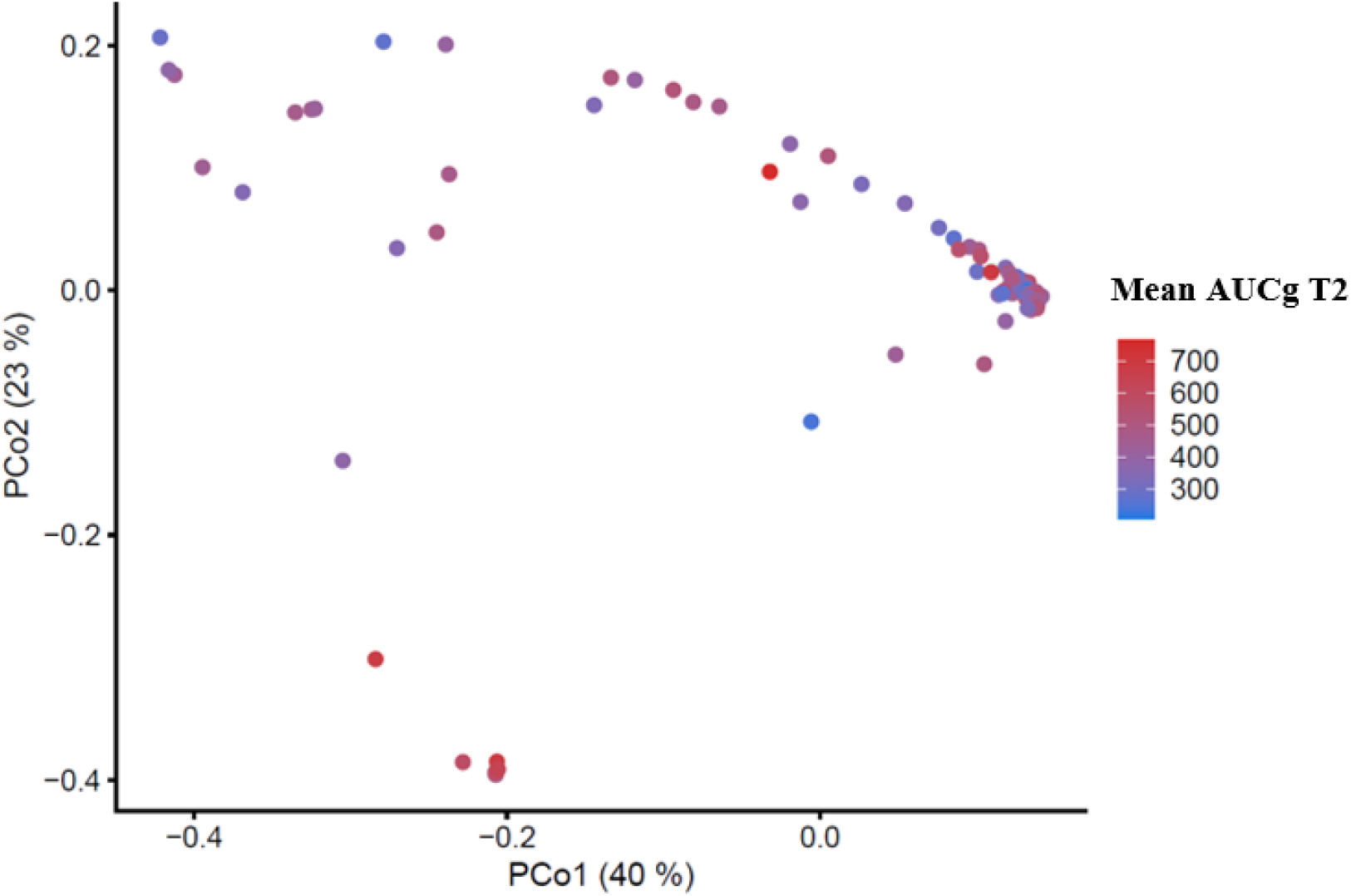
Principal coordinates analysis based on weighted UniFrac distances among meconium samples (*N* = 95), calculated based on the ASV abundances.

### 3.3 Associations between maternal prenatal emotional complaints and the neonatal meconium microbiome

In a first step, we analysed all covariates in relation to the outcome, but found no significant associations between any of the proposed covariates and the microbial composition (p >0.05). Therefore, no covariates were included in the associative analyses.

Results of Kendall rank correlations (see Table 2) revealed that lower alpha diversity of the meconium microbiome was significantly associated with increased maternal prenatal depressive symptoms in late gestation (T1). More specifically, there was a significant association with the Shannon diversity index (τ = -0.149, *p* = 0.041*), which accounts not only for the numbers of ASVs observed in a sample but also for the abundance evenness of the ASVs [66]. In addition, beta diversity was calculated using Bray-Curtis dissimilarity and weighted UniFrac distances, but yielded null findings (Bray-Curtis dissimilarity Pr(>F) = 0.828, weighted UniFrac Pr(>F) = 0.788). Moreover, further calculations revealed no significant findings between prenatal psychological markers (PSS-10, STAI-S) and alpha and beta diversity (Table 2). We further examined the association between the psychological parameters at T1 (EPDS, PSS-10, STAI-S) and the individual taxa from class to species level. However, after using the Benjamini-Hochberg (BH) method to control for FDR [59] at a level of 10%, no correlation with taxon abundances remained significant.

**Table 2.** Alpha and beta diversity associations with psychological parameters (*N* = 95)

| Category | Item | Psychological Variable (T1) | Kendall's tau | p-value |
| --- | --- | --- | --- | --- |
| Alpha Diversity | Shannon diversity index | EPDS | -0.149 | 0.041 * |
| Alpha Diversity | Richness | EPDS | -0.116 | n.s |
| Alpha Diversity | Shannon diversity index | PSS-10 | -0.116 | n.s |
| Alpha Diversity | Richness | PSS-10 | -0.068 | n.s |
| Alpha Diversity | Shannon diversity index | STAI-S | -0.034 | n.s |
| Alpha Diversity | Richness | STAI-S | -0.046 | n.s |

| Category | Item | Psychological Variable (T1) | Sum of Squares | Pr (>F) |
| --- | --- | --- | --- | --- |
| Beta Diversity | Bray-Curtis dissimilarity | EPDS | 0.229 | n.s |
| Beta Diversity | Weighted UniFrac | EPDS | 0.114 | n.s |
| Beta Diversity | Bray-Curtis dissimilarity | PSS-10 | 0.239 | n.s |
| Beta Diversity | Weighted UniFrac | PSS-10 | 0.123 | n.s |
| Beta Diversity | Bray-Curtis dissimilarity | STAI-S | 0.256 | n.s |
| Beta Diversity | Weighted UniFrac | STAI-S | 0.112 | n.s |

Note: $N$ = number of samples; $p$ -value = statistical significance, \* $p < .05$ ; \*\* $p < .01$ ; $Pr(>F)$ = $p$ -value associated with the F statistic; EPDS = Edinburgh Postnatal Depression Scale; PSS-10 = Perceived Stress Scale; STAI-S = State-Anxiety Scale; AUCg = area under the curve with respect to the ground

**Table 3.** Associations of alpha and beta diversity (*N* = 95) and specific bacteria (*N* = 100) with biological stress parameters

| Category | Item | Cortisol | Kendall's tau | p-value |
| --- | --- | --- | --- | --- |
| Alpha Diversity | Shannon diversity index | Mean AUCg (T1) | - 0.078 | n.s |
| Alpha Diversity | Richness | Mean AUCg (T1) | 0.032 | n.s |
| Alpha Diversity | Shannon diversity index | Mean total decline (T1) | 0.002 | n.s |
| Alpha Diversity | Richness | Mean total decline (T1) | - 0.081 | n.s |
| Alpha Diversity | Shannon diversity index | Mean AUCg (T2) | - 0.090 | n.s |
| Alpha Diversity | Richness | Mean AUCg (T2) | - 0.033 | n.s |
| Alpha Diversity | Shannon diversity index | Mean total decline (T2) | 0.096 | n.s |
| Alpha Diversity | Richness | Mean total decline (T2) | 0.073 | n.s |
| Category | Item | Cortisol | Sum of Squares | Pr(>F) |
| Beta Diversity | Bray-Curtis dissimilarity | Mean AUCg (T1) | 0.200 | n.s |
| Beta Diversity | Weighted UniFrac | Mean AUCg (T1) | 0.076 | n.s |
| Beta Diversity | Bray-Curtis dissimilarity | Mean total decline (T1) | 0.233 | n.s |
| Beta Diversity | Weighted UniFrac | Mean total decline (T1) | 0.109 | n.s |
| Beta Diversity | Bray-Curtis dissimilarity | Mean AUCg (T2) | 1.099 | n.s |
| Beta Diversity | Weighted UniFrac | Mean AUCg (T2) | 0.666 | 0.003* |
| Beta Diversity | Bray-Curtis dissimilarity | Mean total decline (T2) | 0.444 | n.s |
| Beta Diversity | Weighted UniFrac | Mean total decline (T2) | 0.251 | n.s |
| Category | Item | Cortisol | Kendall's tau | p-value |
| Species | Brevundimonas naejangsanensis | Mean total decline | 0.270 | 0.0008** |
| Species | Pseudomonas_E | Mean total decline | 0.248 | 0.0006** |

Note: $N$ = number of samples; $p$ -value = statistical significance, \* $p < .05$ ; \*\* $p < .01$ ; $Pr(>F)$ = $p$ -value associated with the F statistic; AUCg = area under the curve with respect to the ground

### 3.4 Associations between prenatal maternal biological stress and the neonatal meconium microbiome

An overview of the analysis is provided in table 3. Results from the dbRDA revealed that the mean AUCg at T2 was significantly related to the beta diversity of the meconium samples (weighted UniFrac Pr(>F) = 0.003*). For a visual interpretation of this association, see Figure 4. No other significant associations were found between salivary cortisol and diversity markers (*p* > .005). We further explored the relationship between cortisol and each taxon from class to species level. After using the Benjamini-Hochberg (BH) method to control for the FDR [59], the mean cortisol total decline at T2 was associated with the species of *Brevundimonas naejangsanensis* and *Pseudomonas_E* (*p* < 0.001**).

### 3.5 Associations between maternal biological stress and emotional complaints

At T1, the EPDS, PSS-10 and STAI-S were not significantly correlated with the AUCg or the mean cortisol total decline (*p* > .05). Similarly, the EPDS, PSS-10 and STAI-S at T1 were not significantly related to the AUCg and the mean cortisol total decline at T2 (*p* >

.05). We further analysed the correlations between cortisol parameters at T1 and T2, and found significant correlations for AUCg (rho = - .419, *p* < .001**) and for mean cortisol total decline between T1 and T2 (rho = .507, *p* < .001**).

## 4 Discussion

In this study following women from late pregnancy until early postpartum, we investigated whether maternal prenatal anxiety, depressive symptoms, and self-reported distress as well as biological stress alter the bacterial composition of the neonate’s first stool discharge after controlling for multiple confounders. Three main findings emerged: Higher maternal depressive symptoms at 34-36 weeks gestation were associated with lower alpha diversity of the meconium microbiome; elevated mean cortisol total decline was associated with the infant’s gut *Proteobacteria*; and higher AUCg was related to higher gut beta diversity in the infant.

The first finding suggests that low gut microbial alpha diversity in neonates is associated with prenatal depressive levels. Similar to previous findings, perinatal anxiety and self-reported distress were not significant predictors of early gut microbiota [30]. There is accumulating evidence that prenatal depression disrupts the healthy development of offspring gut microbiota [67, 68]. Data have confirmed that some species that reach the foetal GI tract prenatally can be present in the infant gut throughout the first year of life [35], highlighting the potential long-term detrimental effects of altered early gut microbiome. Low alpha diversity in infancy may play a role in susceptibility to adulthood diseases and is linked to negative health outcomes in childhood, including asthma [69], type 1 diabetes [70], eczema [71], and skin sensitisation [72]. On the other hand, increased alpha diversity in infants represents a more mature microbial community [73] and higher protection against diseases, and is a predictor of rapid infant growth and functional changes in the gut microbiome [74]. Notably, preliminary findings have revealed a correlation between lower early life alpha diversity and better cognitive development at age two years [73], suggesting that the interpretation of early microbiota alpha diversity in infants is not straightforward. The lower bacterial diversity reported here is likely caused by intrauterine transfer of suboptimal maternal microbiome, which may reflect psychopathological processes occurring during pregnancy [68]. Future studies should consider examining prenatal in-utero microbiota characterisation in order to extend our knowledge of potential transfer mechanisms.

The present findings are in line with earlier studies reporting intrauterine microbial colonisation of meconium [30, 75, 76], as our participants showed an abundance of *Acinetobacter, Pseudomonas, Moraxellaceae and Enterobacteriaceae*. Although *Proteobacteria* was the dominant phylum in our sample, it was not correlated with maternal emotional complaints, but did show a correlation with mean cortisol total decline. This finding partially corresponds to the studies by Zijlmans et al. (2015) [9], Rodriguez et al. (2021) [68] and Wei et al. (2022) [43], who reported that increased maternal stress (both self-reported and measured by cortisol), depression and emotional complaints were related to a higher abundance in *Proteobacteria* in meconium microbiome composition. *Proteobacteria* is an opportunistic gram-negative phylum harbouring lipopolysaccharides in their outer membranes, which is a virulence factor influencing the development of psychopathology, such as anxiety and depression [77]. It is unclear why we found that this phylum was significantly associated with cortisol mean total decline but not with maternal emotional complaints. We believe that this may be due to the low levels of self-reported emotional complaints in our study, while the samples in previous studies [9, 43] showed higher or clinical levels of maternal psychological problems. Whereas cortisol mean total decline levels in late pregnancy were related to *Proteobacteria* in the present study, prenatal depressive symptoms were more detrimental to infant gut microbial diversity than the colonisation with any particular bacteria. This might be viewed as an interesting indicator that the alpha diversity in the meconium microbiome is more sensitive to various depression levels than specific taxa, although this remains to be investigated further.

Our last finding is novel. To the best of our knowledge, there is no previous human study suggesting that increased maternal prenatal biological stress, as measured by cortisol AUCg, is linked to an increase in gut microbial beta diversity in neonates. Beta diversity quantifies inter-individual diversity and measures differences in microbial composition, thus representing a greater maturation of the newborn gut microbiota [74]. The only two previous studies [9, 78] to have assessed maternal prenatal biological stress (e.g. cortisol) in relation to infant faecal microbiota did not find any associations between cortisol and microbiota beta diversity. Differences in the statistical approach used to analyse cortisol data, cortisol assessment tissue, as well as faecal microbiota sampling time and characterisation, preclude direct comparisons between our study and these early studies.

Certain prenatal factors which may contribute to the early colonisation of the gut microbiome showed no associations with our microbiome data. A key advantage of the present study lies in the exclusion of certain confounders and the statistical control of relevant variables such as maternal age, education, parity, maternal BMI, gestational age, mode of delivery, newborn length and weight, and meconium collection site. This provided substantial coverage of the environmental determinants of meconium microbiota. Participants were excluded from the study if they received antibiotic treatment over the course of pregnancy, but women who underwent an unplanned Caesarean section and received a dose of antibiotics during delivery were retained in the analyses. This did not change the overall study conclusions. As expected, the birth mode did not significantly affect microbiota, as gastrointestinal tract colonisation is mostly derived from the intrauterine environment [79]. A recent study compared meconium microbiota to amniotic, vaginal, and oral cavity microbiotas, and found the greatest overlap between meconium and amniotic datasets [80]. Importantly, all meconium samples were obtained prior to feeding; thus, infant diet should have exerted no effect on microbial biomarkers.

Our study offers a number of strengths, including following women from late pregnancy to early postpartum, the early assessment of newborn meconium, the measurement of psychological emotional complaints and biological markers of stress, and the collection of detailed information about potentially important confounding variables. However, some clear limitations should also be mentioned. The sample was generally well educated and not economically diverse. Moreover, the study may have benefited from additional assessments of both biological and self-reported measures at all time points. Future studies combining longitudinal time-series profiling with bacterial strain-level resolution sequencing may be better placed to identify the dynamics of bacterial strains by the level of maternal emotional complaints. Additionally, the study does not allow for conclusions regarding causative biological pathways through which psychobiological measures influence the neonatal meconium; however, this was not our aim. Our data provide a novel insight into the impact that maternal depressive symptoms and cortisol in late pregnancy may have on the neonate’s meconium microbial community.

In sum, higher maternal prenatal depressive levels and cortisol were significantly associated with the neonate’s microbiota in a cross-sectional human cohort. Furthermore, the altered alpha diversity pattern and increased *Proteobacteria* suggests that children should be followed up, as both have been linked to negative health outcomes. Our novel finding warrants further research and replication. The present results shed more light on the meconium microbiome composition, helping to elucidate how the maternal prenatal environment interacts with the initial bacterial composition of the neonate. These results corroborate some previous findings and suggest hypotheses for future studies.

## Abbreviations

ASV: amplicon sequence variant
AUCg: area under the curve with respect to the ground
BH: Benjamini-Hochberg
dbRDA: distance-based redundancy analysis
EPDS: Edinburgh Postnatal Depression Scale
FDR: False discovery rate
GAS: Geburts-Angst-Skala
GHQ-12: General Health Questionnaire
HPA: axis hypothalamic-pituitary-adrenal axis
NuPDQ: Prenatal Distress Questionnaire
PSS-10: Perceived Stress Scale
STAI-S: State-Trate Anxiety Inventory, state scale

## Acknowledgements

We are grateful to the Research Group for Food Perception of the ZHAW School of Life Sciences and to the Swiss Federal Food Safety and Veterinary Office for their support. We would like to thank Annika Schneider, Eileen Stephan, Nathalie Appenzeller, Rhea Sarah Gesù, Tamara Grossrieder, Tamara Lovrinovic, Seraina Weber, Stéphanie Loeffel, Simon Hühni and Firouzeh Farahmand for their research assistantship. We also would like to thank Rausch AG, Medela Schweiz AG, MAM Baby AG and Bimbosan which freely provided gift sets for study participants. Finally, we wish to express our gratitude to all women who took part in this study.

## Data availability statement

Raw sequencing data for all 100 samples described in this project have been deposited in The Open Science Framework (OSF) under osf.io/r687v. The data that support the findings of this specific study are available from the corresponding author upon reasonable request.

